# Antibiotic resistance genes and class 1 integron: Evidence of fecal pollution as a major driver for their abundance in water and sediments impacted by metal contamination and wastewater in the Andean region of Bolivia

**DOI:** 10.1101/2020.03.23.003350

**Authors:** Jorge Agramont, Sergio Gutierrez-Cortez, Enrique Joffré, Åsa Sjöling, Carla Calderon Toledo

**Affiliations:** Environmental Microbiology Unit. Institute of Molecular Biology and Biotechnology, Universidad Mayor de San Andrés, La Paz, Bolivia; Department of Microbiology, Tumor and Cell Biology, Karolinska Institutet, Stockholm, Sweden; Centre for Translational Microbiome Research, Karolinska Institutet, Stockholm, Sweden

## Abstract

Water and sediment samples affected by mining activities were collected from three lakes in Bolivia, the pristine Andean lake Pata Khota, the Milluni Chico lake directly impacted by acid mine drainage, and the Uru-Uru lake located close to Oruro city and highly polluted by mining activities and human wastewater discharges. Physicochemical parameters, including metal compositions, were analyzed in water and sediment samples. Antibiotic resistance genes (ARGs), were screened for, and verified by quantitative PCR together with the mobile element class 1 integron (*intl1*) as well as crAssphage, a marker of human fecal pollution. The gene *intl1* showed a positive correlation with *sul1, sul2, tetA* and *blaOXA-2*. CrAssphage was only detected in Uru-Uru lake and its tributaries and significantly higher abundance of ARGs were found in these sites. Multivariate analysis showed that crAssphage abundance, electrical conductivity and pH were positively correlated with higher levels of *intl1* and ARGs. Taken together our results suggest that fecal pollution is the major driver of higher ARGs and *intl1* in wastewater and mining contaminated environments.

## Introduction

Antibiotic resistance is considered a major threat to human health worldwide (1). Antibiotic resistance genes (ARGs) have been classified as emergent contaminants, with a significant impact in aquatic environments due to the possibility to be acquired by pathogens, which could lead to public health issues (2). Novel rearrangements of ARGs and mobile genetic elements (MGEs), that favor their dissemination, are considered xenogenetic pollutants. These elements can be incorporated and replicated in environmental microorganisms, thereby increasing their concentration (3).

It has been reported that anthropogenic activities cause pollution of aquatic environments with ARGs and MGEs (4). Wastewater discharges cause co-occurrence of MGEs and different ARGs in water and sediments (5). At a continental scale, ARGs in sediments are strongly correlated with MGEs and antibiotic residues (6). Recently, it has been observed that microorganisms living in aquatic microbial communities that came from wastewater were able to transfer ARGs via horizontal gene transfer (HGT) after exposure to low levels of antibiotics and biocides (7). Many of the ARGs that can be found in clinical settings have also been found in the environment, suggesting the possibility of movement and dissemination between these two scenarios (8).

Mining activities cause contamination of downstream water with dissolved metals (9) where heavy metals tend to accumulate in sediments (10). It has been suggested that heavy metals can favor selection of ARGs via co-selection, *i.e*. the simultaneous acquisition of both, ARG and metal resistance genes, where the presence of metals constitutes the selective pressure (11). Several studies support this relation. Urban soil samples of Belfast in Northern Ireland, exhibited a pattern of co-occurrence between metals (Zn, Cu, Cd, Co, Ni, Hg, Cr and As) and many ARGs. Moreover, the degree of metal toxicity was positively correlated with the abundance of MGEs, and ARGs (12). Metals, as Cu and Zncan in some cases exert stronger selection pressure over soil microbial communities for the selection of resistant bacteria, even more than specific antibiotics (13). In the Dongying river in China, the levels of Cu and Cr were positively correlated with the abundance of different ARGs (14), whereas both Zn and Pb levels were correlated with the abundance of erythromycin resistance genes in wastewater treatment plants (15). These data suggest that, aquatic environments are important for the ecology and evolution of ARGs. In particular, water bodies can be hotspots for the evolution of ARGs due to the convergence of antibiotics, microorganisms from different sources, biocides and heavy metals (16), generating a scenario that favors the emergence, persistence and dissemination of ARGs (17).

Levels of fecal pollution are not frequently considered in the analysis of selection and dissemination of ARGs (18). The incorporation of a molecular marker of human fecal pollution can help us to disentangle the accumulation of ARGs due to fecal bacterial discharges, and the ARGs selection and dissemination caused by other environmental contaminants (18). CrAssphage, most probably infect *Bacteroides* and *Prevotella* bacteria in the human gut (19). The gene KP06_gp31 that belongs to CrAssphage is highly abundant in aquatic environments contaminated by human feces, while it is less abundant in aquatic environments polluted by feces of other animals (20); thus, the CrAssphage can be considered a marker for human fecal pollution.

Mining activities, which are known to have a great impact on water resources (21, 22), have a long history in the Bolivian Andean region. This region that, includes La Paz, El Alto and Oruro cities, is going through a water scarcity process (23) as a consequence of climate change effect on glaciers (23). Furthermore, urban wastewater is directly released into the environment, polluting water with enteric pathogens and resistant bacteria (24, 25). To our knowledge, there are no previous studies in the region that consider the problem of accumulation of emergent contaminants such as ARGs, MGEs in water and sediments, of mining impacted sites and water reservoirs, taking into consideration the effects of wastewater discharges through the use of a human fecal pollution molecular marker.

The aim of this work was to analyze the influence of metal pollution, and human fecal discharges on the abundance of different ARGs and the class 1 Integron (*intl1*), in water and sediments samples from a pristine, metal polluted and wastewater-mining contaminated lakes, in order to explore the contributions of metals and fecal discharges in the abundance of ARGs. Our results showed an increased abundance of class 1 integron and ARGs in correlation with the levels of fecal pollution. Fecal polluted sites presented significantly higher levels of *intl1* and ARGs. Moreover, multivariate analysis showed that AGRs and *intl1* abundances were positively related with the abundance of crAssphage, and physicochemical parameters (pH and EC), suggesting that fecal bacterial contributions are the main responsible for the increased abundance of ARGs into the environment.

## Materials and methods

### Sampling sites

#### The Milluni valley Lakes

Milluni valley is located in the Andean region of Bolivia, in the Department of La Paz, 20 Km from La Paz city in the Cordillera Real. It is a glacial valley at the foot of the Huayna Potosi mountain. The valley has four lakes: Pata Khota (4670 masl), Jankho Khota (4575 masl), Milluni Chico (4540 masl), and Milluni Grande (4530 masl) the biggest one, with a surface of 2.37 Km^2^ and 4 m depth. Milluni Grande has a dam that captures water that supplies water to the Puchucollo drinking water treatment plant, then water is distributed to the cities of La Paz and El Alto.

Mining activities were performed in the valley until 1990 by the company COMSUR, water from the surrounding lakes was used in the mining activities. Acid mining drainage (AMD) were discharged directly in the Milluni Chico lake, also contaminating the downstream lake of Milluni Grande. As a consequence, these two lakes acquired an extremely acid pH that favored the mobility of metals (Cd, Zn, As, Cu, Ni, Pb, Sn) in water and sediments (26). In contrast, the first lake Pata Khota, is fed by water proceeding from the melting of Huayna Potosi mountain. Anthropogenic activities are very limited around this site, and the lake is considered an ecologically intact environment (26, 27).

#### Uru Uru Lake

The Uru Uru Lake (3686 masl), situated in the department of Oruro, in the central part of the Altiplano in Bolivia, is an artificial shallow lake 8 Km south of Oruro city. The lake is characterized by an alkaline pH (8,3 ± 0,6) with a strong buffering capacity (28). The Tagarete channel receives and drains the wastewater from Oruro city towards the northern part of the lake. The north-east part of Uru Uru lake receives water discharges from San Jose Mine and the Vinto smelting plant (28). On the other hand, the Desaguadero River that comes from the Titicaca Lake drain the discharges of Kori Kollo and Kori Chaca meromictic lakes (once open pit gold mines) into the north-west part of Uru Uru (29). Previous studies in Uru Uru, reported that the contribution of both, wastewater and mining residues increase the electrical conductivity (EC) and the concentration of certain metals and metalloids such as: Hg, Fe, Mn, W, and Sb. (28)

### Sample collection and processing

Milluni samples were collected during the dry season in July 2016. Three points were randomly selected in Milluni Chico (MC) and Pata Khota (PK) lakes (Fig 1). Temperature, electrical conductivity (EC) and pH were measured directly on water (Oakton Instruments, Vernon Hills). Duplicate superficial sediment samples were collected in sterile 50 mL centrifuge tubes, for both DNA extraction and metal quantification. Samples were immediately labeled and stored at 4°C with cold packs and rapidly transported to the laboratory where they were stored at - 70°C until their analysis.

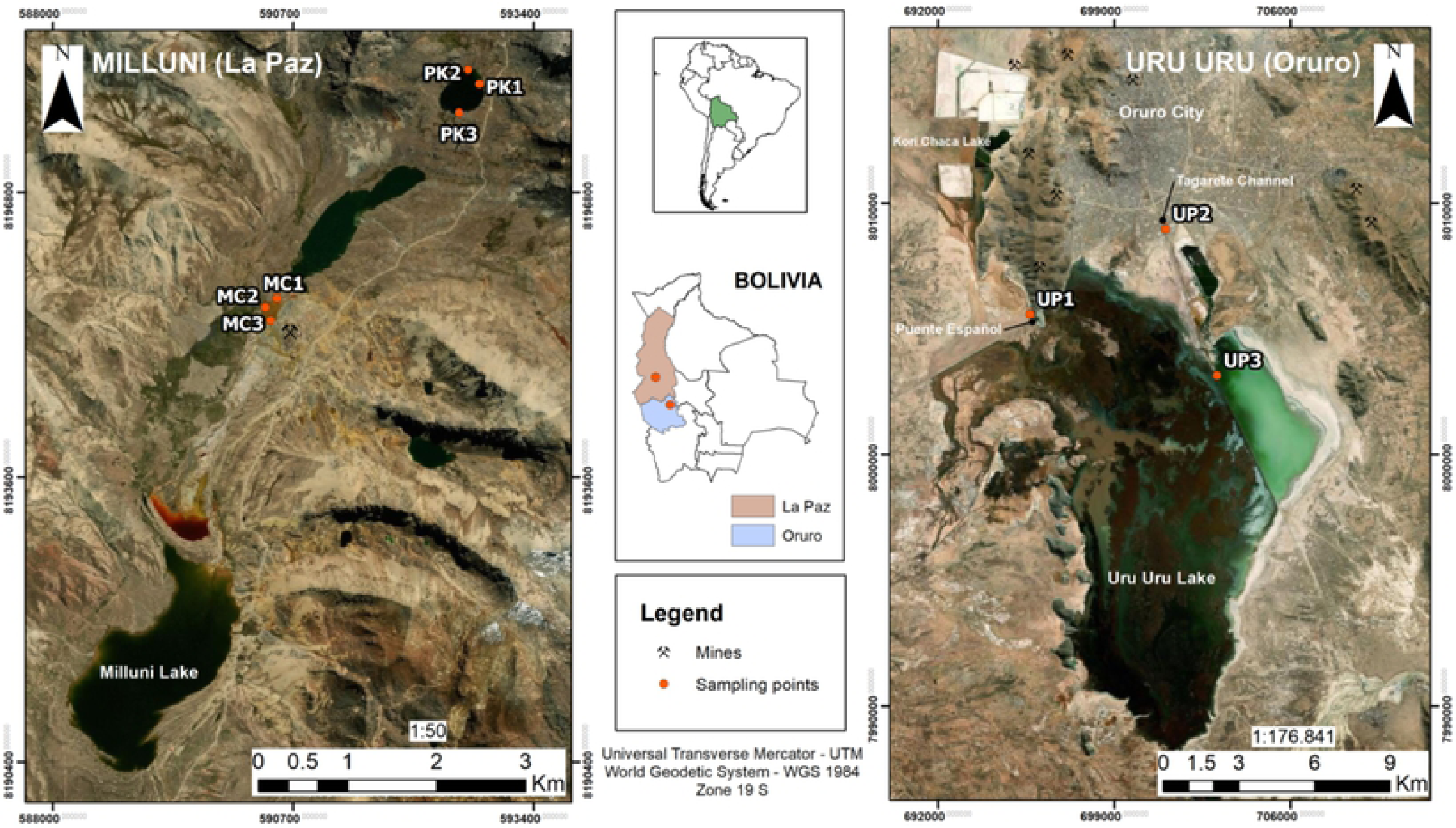

Samples of Uru Uru and its tributaries were collected at the beginning of the rainy season (November 2018). Three different points were considered: **(1)** UP1: the channel that discharges the water of the meromictic lake Kori Chaca into the northwest part of Uru Uru lake. Agricultural activities are performed around this channel, and wastewater discharges were previously reported (30); **(2)** UP2: the Tagarete channel that carries untreated wastewater discharges from Oruro city; and **(3)** UP3: located in the north east part of the lake, where Tagarete’s discharges drain.

Sediment samples were collected in triplicate. Parameters were recorded as described for Milluni. Surface sediment samples were collected in sterile 50 mL centrifuge tubes, using a Core Sampling Device. The samples were divided into two fractions, one for DNA extraction and the other for metal quantification.

Samples were kept at 4°C with cold packs, and transported to the laboratory in La Paz city, were they were rapidly stored at −70°C until their analysis. Surface water samples from UP1, UP2 and UP3 were collected in triplicate, filtered (300 mL) through 45 µm nitrocellulose filter membranes (Sigma-Aldrich), and the filters were immediately stored at −70°C until their analysis.

### Quantification of metals

Six elements were quantified in the sediment and acidified water samples: Cu, Zn, Pb, Ni, Cd, and As. All the analyses were performed as previously described (30). The measurement was performed using Inductively coupled plasma mass spectrometry. The quantification was performed at the *Laboratorio de Calidad Ambiental* (LCA), Universidad Mayor de San Andres.

### DNA extraction

DNA was extracted from sediments using PowerSoil DNA isolation kit (Qiagen, Germany). A prewashing step was performed using solution S0 (0.1 M EDTA, 0.1 M Tris (pH 8.0), 1.5 M NaCl, 0.1 M NaH_2_PO_4_, and Na_2_HPO_4_) (31), due to the acidity of some samples and the presence of heavy metals. Briefly, 300 mg of sediments were washed with 1,5 mL of solution S0 overnight, in a horizontal shaker at 180 rpm at 4°C, the sediment was recovered by centrifugation at 12 000 × g for 5 min and repeatedly washed with S0 until the supernatant end up clear. After washing, the sediments were transferred to PowerBead Tubes (Qiagen, Germany) and the extraction proceeded as described by the manufacturer’s instructions.

Additionally, a quarter of the filtered water samples was used for DNA extraction using the PowerSoil DNA isolation kit (Qiagen, Germany). The filter was transferred into the *PowerBead* Tubes (Qiagen, Germany) to proceed with the DNA extraction according to the manufacturer’s protocol. DNA concentration was measured using Qubit® dsDNA HS (Invitrogen, Oregon USA).

### Quantitative PCR

Selection of ARGs for the analysis was performed as follows: The Antibiotic Resistance Genes Microbial DNA qPCR arrays (Qiagen, USA) were used to screen for the presence of ARGs. The array consists of 96 well plates with predispensed primers for 85 different ARGs (S1 Table) conferring resistance to antibiotics commonly used in clinical settings. Twelve positive ARGs (CT < 39) were selected to perform the assays of absolute quantification. Additionally, sulfonamides resistance genes (*Sul1* and *Sul2*) were included in the analysis as previous reports point at their presence in Milluni (32).

Standard curves for the absolute quantification of target genes were constructed using a plasmid as a template (S1 Fig). This plasmid was engineered by the insertion of the PCR assembled products of 14 ARGs (β-lactams [*acc-3, bla*_I*MP-2*_, *bla*_IMP-5_, *bla*_IMP-12_ and *bla*_OXA-2_], macrolide-lincosamide-streptogramin B (*msrA*), methicillin (*mecA*), quinolones (*qnrB1, qnrB5* and *qnrS1*), tetracycline (*tetA* and *tetB*), and sulfonamides (*sul1* and *sul2*)), the class 1 integron gene (*intl1*) and the KP06_gp31 gene of the crAssphage, into the *Xba*I restriction site at the MCS of the pUC57 vector. The assembled sequence was synthesized and inserted by GenScript (Genscript, USA). Reference sequences for the ARGs were obtained from The Comprehensive Antibiotic Resistance Database (CARD) (33) and primers (Table 1) were designed using Primer-BLAST (NCBI) (34). A six point calibration curve was generated using serial dilution from 10^6^ to 10^1^ copies of the plasmid. The 16S rRNA housekeeping gene was used for the normalization of the absolute quantification of ARGs.

**Table 1.**
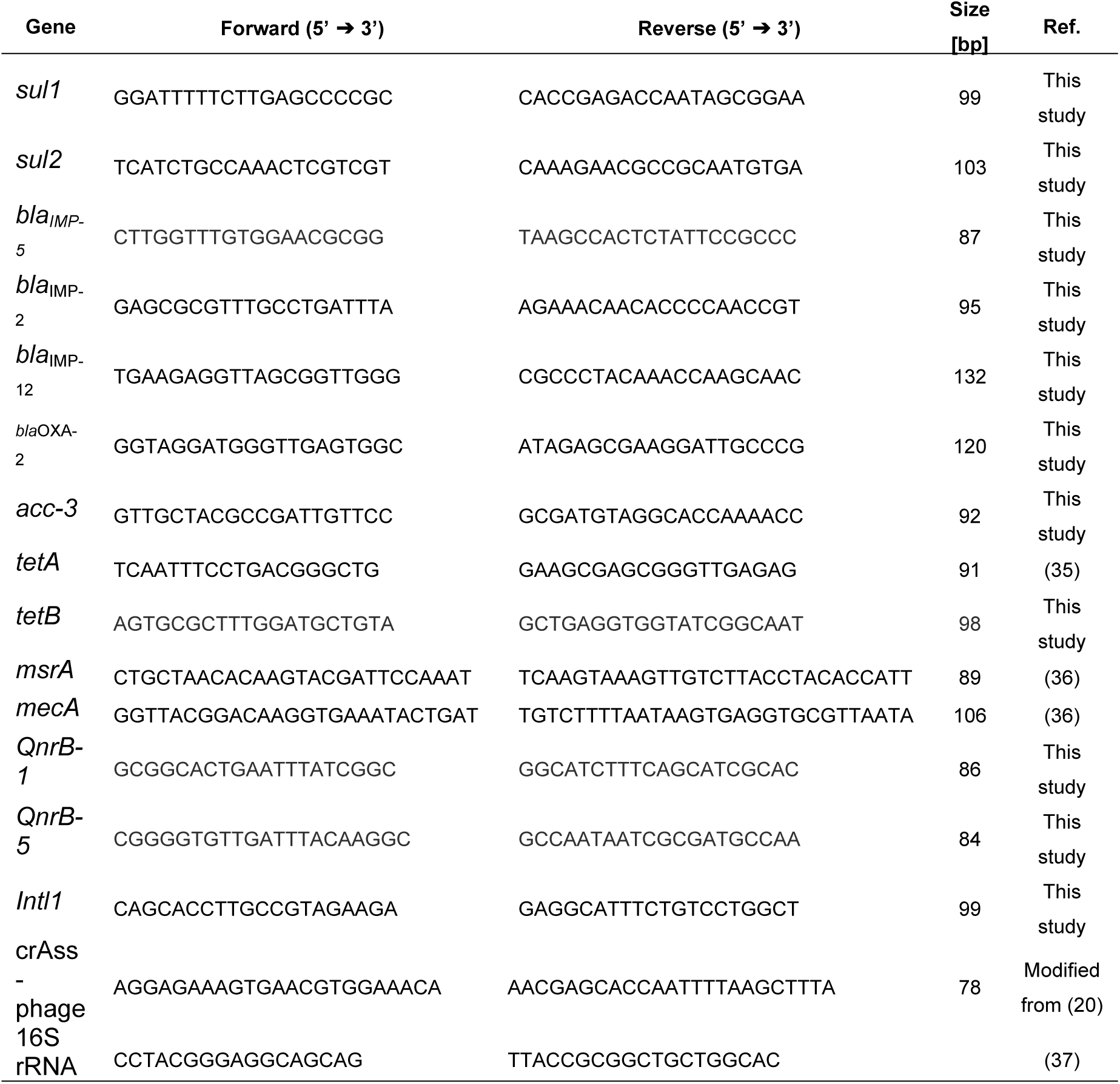
Primers used for quantitative PCR experiments.

For qPCR experiments, the reaction mix consisted in 12.5 μl of Power SYBR® Green PCR Master Mix (Applied Biosystems), BSA 0.1 μg/μl (New England Biolabs), 10 pmol of each primer, 2 μl of DNA, in a final volume of 25 μl. All qPCR experiments were performed in a LightCycler® 480 Instrument II (Roche Molecular Diagnosis), with the following conditions: an initial denaturalization cycle at 95 °C during 5 minutes, followed by 45 cycles of denaturalization at 95°C per 15 seconds, and amplification at 60°C per 30 seconds. A melting curve was performed. All the qPCR experiments were performed at the Centre for Translational Microbiome Research (CTMR) Karolinska Institutet.

The absolute abundance of ARGs and *intl1* gene, was normalized to the absolute abundance of the 16S rRNA gene, as described before (6). The normalized abundance was corrected, in order to express the abundance of genes per bacteria, assuming that the average number of 16S rRNA genes per bacteria is four (6). Absolute abundance for the KP06_gp31 gene (crAssphage) was considered for all the analysis.

### Statistical Analysis

All statistical analysis were performed in the Software R 3.6.1. ANOVA test was performed for the account of statistically significant differences of physicochemical parameters between sites. Previously, the distribution of residues and the homogeneity of variance was analyzed using ‘*qqPlot*’ and diagrams of dispersion respectively. When the data did not fit with the residue distribution and/or the homogeneity of variance, the *‘Box-cox*’ function was used to choose an appropriated transformation. When ANOVA test was significant, pairwise comparisons among means via Post-hoc Tukey was performed to find out differences among sample sites. The package ‘*multcomp*’ (version 1.4-10) was used for multiple comparisons. Pearson correlations were performed in order to evaluate correlation among physicochemical parameters and among quantified genes.

A heat map with hierarchical clustering ordination was performed to present the normalized abundance of the quantified genes, using the R package gplots (version 3.0.1.1).

A Principal Component Analysis (PCA) was performed in order to examine the relationship between the quantified genes, the contributions of fecal discharges, the physicochemical variables and the measured elements. Sampling points were ordered in function of the normalized abundance of ARGs and *Intl1*. The two axis that explained most of the variation were extracted and a multiparametric linear regression (Lm) was used to relate the ordination of the sampling points along the axis with the environmental variables. The selection of the best regression model was automatically performed using the function ‘*regsubsets*’ (backward and forward) from the package ‘*leaps*’ (version 3.1) in function of R^2^ adjusted values. The package *‘car’* (version 3.0-3) was used to calculate the variance inflation factor (VIF) of the independent variables to avoid multicollinearity. The packages *‘factoextra’* (version 1.0.6), and *‘vegan’* (version 2.5-5) were employed for ordination analysis. Both data sets, the abundance of ARGs, *intl1*, and the explaining variables (EC, pH, metal levels and the fecal pollution marker crAssphage) were logarithmically transformed before the analysis.

## Results

### Physicochemical conditions and metal levels

We first evaluated the physicochemical characteristics and metal concentration on every sampling site. The measurement of EC and pH from water samples was performed *in situ* and before the collection of sediments. As shown in Figure 2a, the shallow water lake of Milluni Chico (MC) presented the lowest pH levels (2.32±0.06 in 3 samples). MC lake showed acid mining drainage (AMD) discharges and intense orange color. Pata Khota (PK) sampling sites, and the Uru Uru lake had pH values close to neutral (Fig 3a). The EC measurements showed that PK samples registered the lowest EC values, while MC and Uru Uru water samples showed a significant ten to twenty-fold increase compared to PK, with the exception of UP1(Fig 2b).

**Fig 2.**
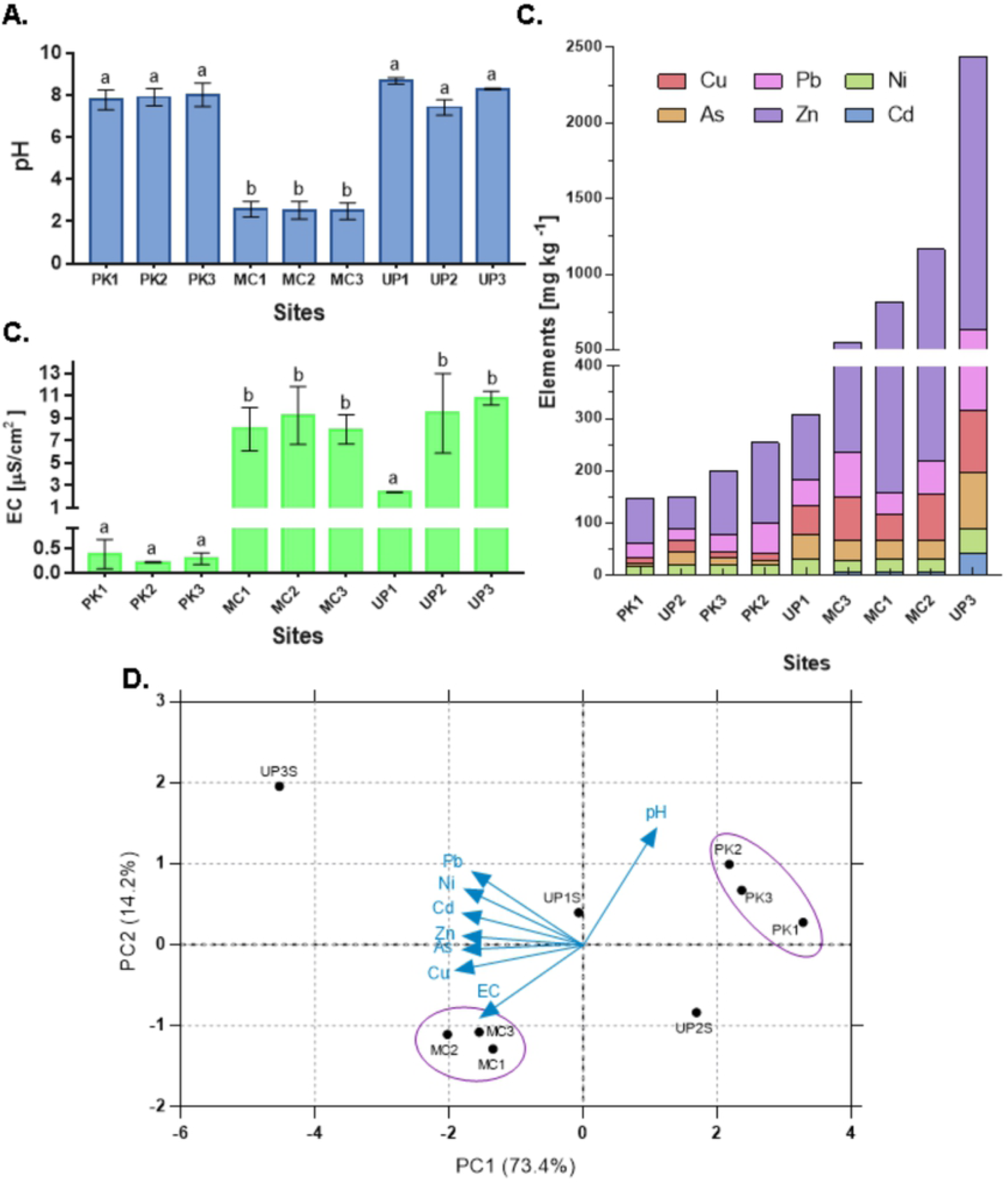
Physicochemical parameters and metal levels. The mean values are shown in bars and standard deviation values are presented as the error-bars. Different letters represent statistically significant differences (p < 0.05), calculated by ANOVA using the Software R 3.6.1. **(A)** pH of the water: MC sampling points, directly impacted by AMD presented the lowest pH. **(B)** Electrical conductivity (EC): PK samples presented the lowest values of EC, whereas wastewater and mining discharges receiving points presented higher values of EC. **(C)** Levels of quantified elements: The levels of metals in sediments are shown in bars with cumulative values. UP3 was the point withthe highest level of all elements, followed by MC and UP1, all of them directly impacted by mining discharges, whereas, UP2 and PKsamples presented the lowest levels of metals. **(D)** Principal Component Analysis (PCA) of physicochemical parameters and metal concentrations of sampling sites: All the sampling points in PK lake presented very similar values for the different environmental parameters, clustering together in the figure. The same was observed for MC sampling points, whereas Uru Uru sampling points did not cluster together, each of the samples presented a clear difference in their metal concentrations.

**Fig 3.**
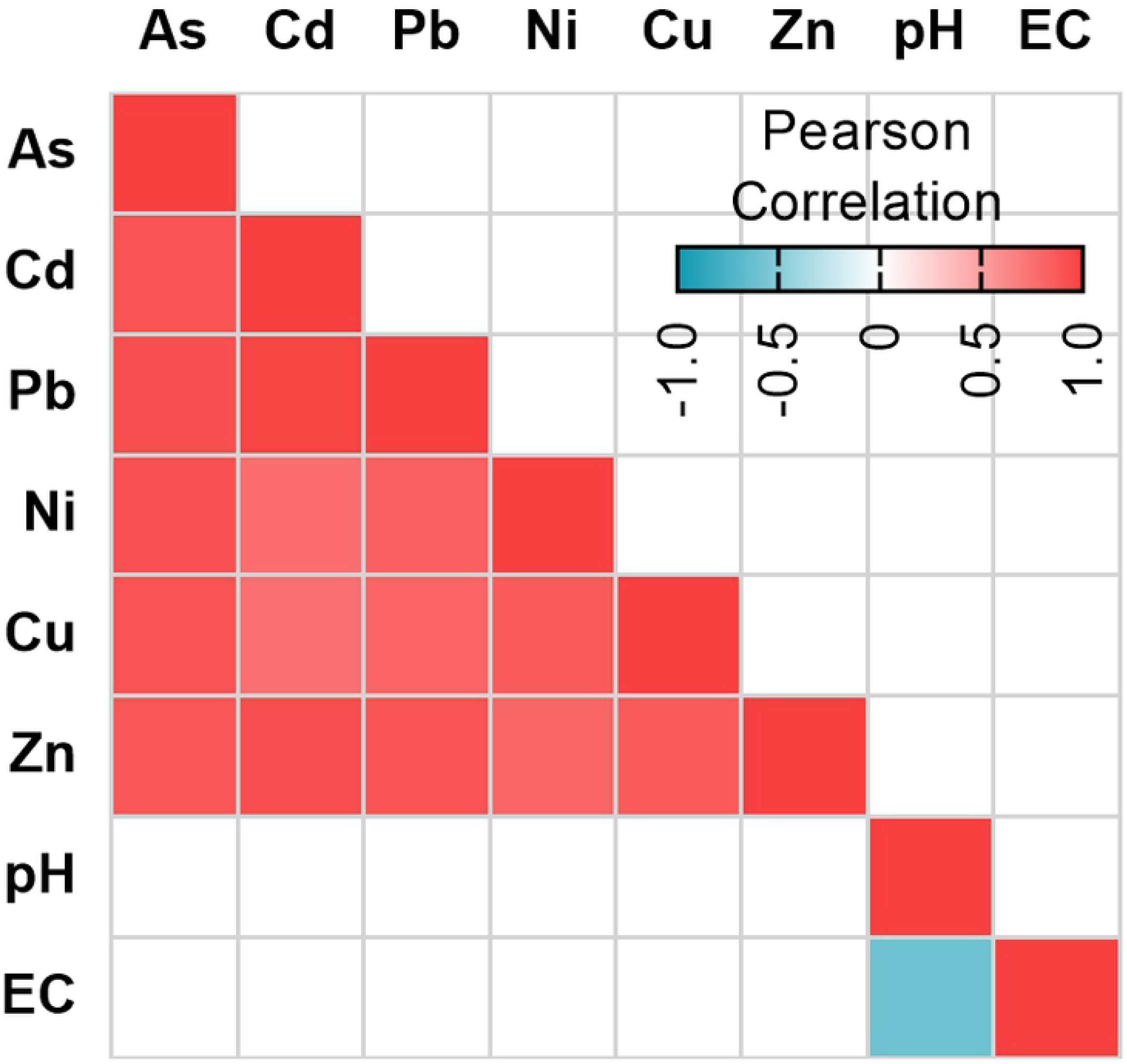
Correlation among metals and physicochemical parameters. The R correlation coefficient is represented in colors as indicated in the legend. Only significant correlations (p<0.05) were included. All elements showed positive correlation among them and with pH. EC and pH were negatively correlated.

Sediments were quantified for the presence of 6 elements (As, Cd, Pb, Ni, Cu, and Zn). The results (Fig 2c) indicated clear differences between sample sites in their metal composition. UP3 (Uru Uru Lake) had the highest concentration of Zn (1811 mg Kg^-1^ of sediment). In comparison with all other samples, statistically significant higher levels of As, Cd, Pb, Ni, and Cu, were found in UP3 except for Cu Cu levels in MC sediments. After UP3, MC samples were the second most abundant for all the elements analyzed, with significantly higher concentrations than UP2, and significantly higher concentrations of As, Cd, Ni, Cu and Zn in relation to PK samples, and also significantly higher than UP1 except for As, Cd, and Pb, levels. UP1 presented higher levels of As, Pb, Ni, and Cu than UP2. Thus, PK samples and UP2 were the sites with the lowest levels of all elements, except for Cu, with significantly higher concentrations in UP2. In summary, all the points directly impacted by mining activities (UP3 followed by MC sites and UP1) contained higher concentrations of As, Cd, Cu and Zn.

A Principal Component Analysis (PCA) was performed to visualize the distribution of the sampling points based on their environmental variables (Fig 2d). All the sampling points in Pata Khota (PK) lake presented very similar values, and were grouped together. Similar trend was observed for Milluni Chico (MC) sampling points. In contrast, Uru Uru sampling points were clearly differentiated by their physicochemical parameters and metal concentrations.

The correlation analysis of all the physicochemical parameters, and metal levels (Fig. 3) showed that metals levels were positively correlated among them. EC and pH showed a negative correlation between them and no significant correlation with other environmental parameters was observed.

### Detection and quantification of ARGs and MGEs

In order to determine and quantify the presence of ARG, we extracted total DNA from the microbial communities present in the sediments and water from the sampled sites. The presence of ARGs was first screened by qPCR using the Microbial DNA Array for Antibiotic Resistance (Qiagen, Hilden, Germany) on PK and MC samples. Based on these results, a plasmid containing fourteen ARGs was designed and constructed. The ARGs sequences inserted included resistance to tetracycline (*tetA, tetB*), β-lactam antibiotics (*bla*_OXA-2_, *bla*_IMP-2_, *bla*_IMP-5_, *bla*_IMP-12_, *acc-3*), methicillin (*mecA*), quinolones (*qnrB1, qnrB5* and *qnrS1*), and macrolide-lincosamide-streptogramin B (*msrA*) sulfamethoxazole (*sul1* and *sul2*). The latter two were not included in the screening arrays but were previously reported in the area (32). In addition the sequences of the bacteriophage crAssphage and Class 1 integron (*intl1*), were inserted in the same plasmid. This plasmid was used to generate standard curves for all qPCR runs and samples analyzed.

The normalized abundance of these genes is shown in Fig 4. The *acc-3* genes were only detected in the Tagarete samples (UP2), both in sediments and water. The Class 1 Integron, together with *bla*_*OXA-2*_ and *sul1* sequences were detected in all the samples. The fecal contamination marker crAssphage was only detected in sediments and water from Uru Uru and its tributaries, except for UP1 in which it was only present in water. The UP2 site that receives wastewater discharges presented the highest crAssphage abundance. In contrast crAssphage was not detected in PK and MC samples. In general, UP2 and UP3 samples were the ones with the highest abundance for the majority of the quantified genes, being *intl1, sul1, sul2* and *bla*_*OXA-2*_ the most abundant. Except for PK3, the hierarchical clustering analysis revealed a distint gene abundance between Milluni basin and Uru Uru sites.

**Fig 4.**
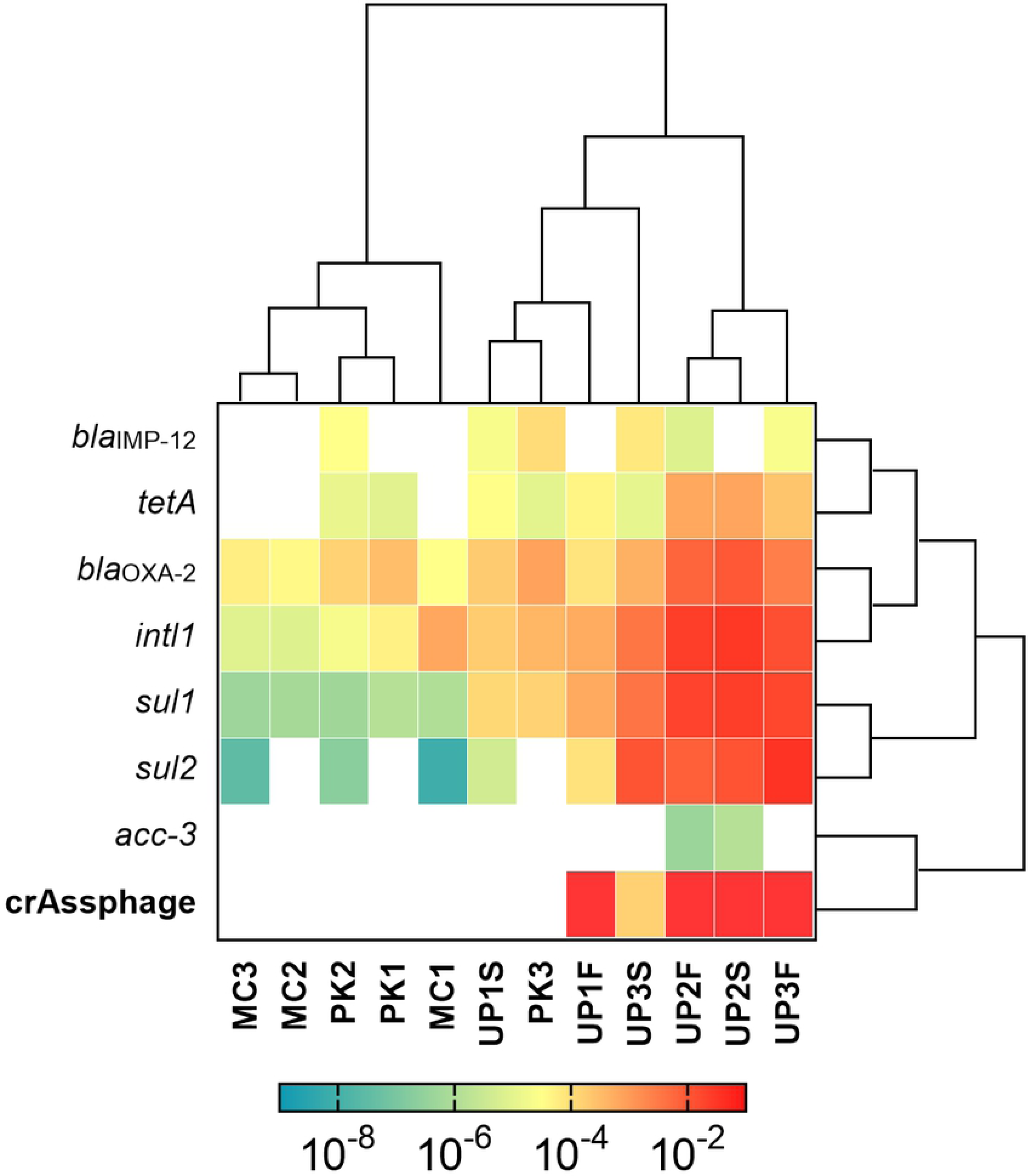
Normalized abundance of ARGs, *intl1* and CrAssphage detected on sediments and water. The abundance values of ARGs and *intl1* were normalized in function of 16S rRNA gene abundance, absolute abundance of crAssphage is included. The data were transformed using Log_(10)_ and represented in a heatmap where, reddish coloration symbolizes a higher abundance. Rows and columns were ordered by similarity with a hierarchical clustering. CrAssphage was detected exclusively in Uru Uru samples, with the exception of UP1 sediments, UP2 sampling sites presented the highest abundance of this gene and were the only ones with *acc3* gene signal. *intl1, sul1*, and *bla*_OXA-2_ were found in all samples, with similar abundance patterns, being most abundant in Uru Uru. S: sediments; W: water.

The correlations of the normalized abundance among all detected genes were evaluated (Fig 5). Class 1 integron was positively correlated with *sul1, sul2, bla*_OXA-2_, and *tetA* genes. Furthermore, these five genes had also positive correlation among them, except *sul2* and *tetA*. The abundance of *tetA* presented an inverse correlation with *bla*_IMP-12_. crAssphage absolute abundance was not correlated with the abundance of any other gene.

**Fig 5.**
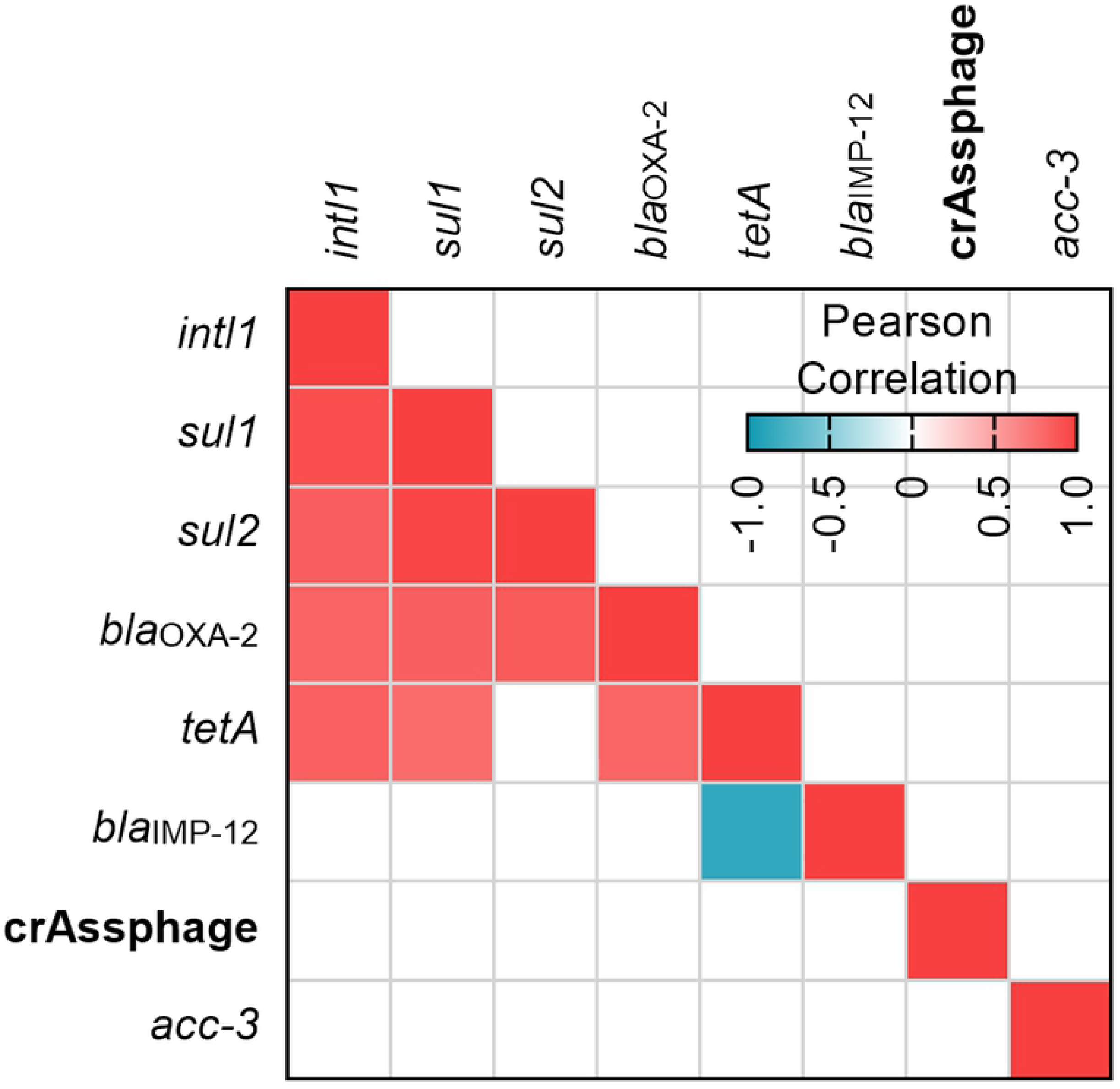
Correlation among the detected genes. R correlation coefficient is represented as a heat map of colors. Only significative correlations (p < 0.05) are depicted. *intl1* abundance positively correlated with the abundance of *sul1, sul2, bla*_OXA-2_ and *tetA* genes, all these genes (with the exception of *tetA* with *sul2*) presented positive correlations among them. The fecal pollution marker did not correlate with any other gene.

### Fecal pollution and antibiotic resistance

To evaluate if fecal pollution contributes to the abundance of *intl1* and ARGs, a one-way ANOVA was performed to compare the abundance of *intl1* between samples with and without the presence of crAssphage. Samples in which crAssphage was detected, presented a significantly higher abundance of *intl1* (p<0.001). Moreover, the abundance of each ARG that positively correlated with *inlt1* (i.e. *sul1, sul2, bla*_OXA-2_, and *tetA*) presented a statistically higher abundance in sampling sites with positive signal for crAssphage (Fig 6).

**Fig 6.**
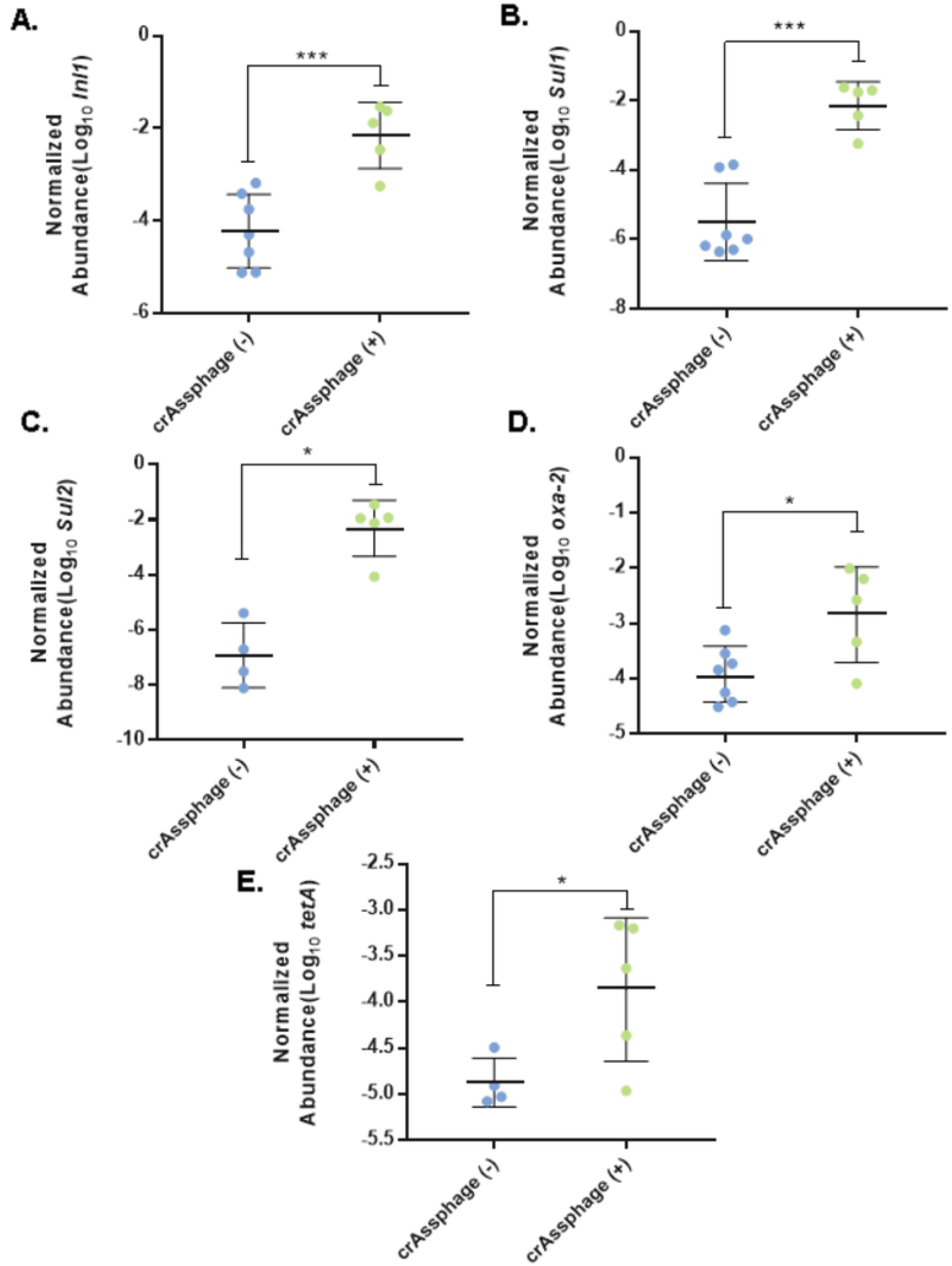
ANOVA of the abundance of ARGs and *intl1* in function to the presence of crAssphage. By comparing the abundance of *intl1* between the samples with detected crAssphage and samples in which the fecal marker was not detected, statistically significant differences were found, being fecal polluted samples the ones with the highest abundance of *intl1* gene. The same pattern was found for *sul1, sul2, bla*_OXA-2_ and *tetA*.

### Relationship between ARG and environmental variables

To evaluate the relationship between the abundance of ARGs, *intl1*, fecal pollution (crAssphage) and the environmental factors (metal levels and physicochemical parameters) a PCA was performed. ARGs and *intl1* abundance were reduced into the first two principal components (PCs) that explained 69.5% and 16.3% of the variation respectively, these two PCs were recovered. A linear model was performed in order to see if along the PCs the ordination of the samples in function of the abundance of genes respond to any environmental variable (Fig 7). The linear regression showed that the PC1 presented a positive linear relation with crAssphage, pH, and EC (Adj. R^2^ = 0.969; F= 50.52; p< 0.05), suggesting that fecal pollution, a neutral pH value, and high EC are the three conditions related with higher abundance of ARGs and *intl1*. No statistically significant relationship was found for the second PC. We could not find any significat association between metal levels and gene abundances.

**Fig 7.**
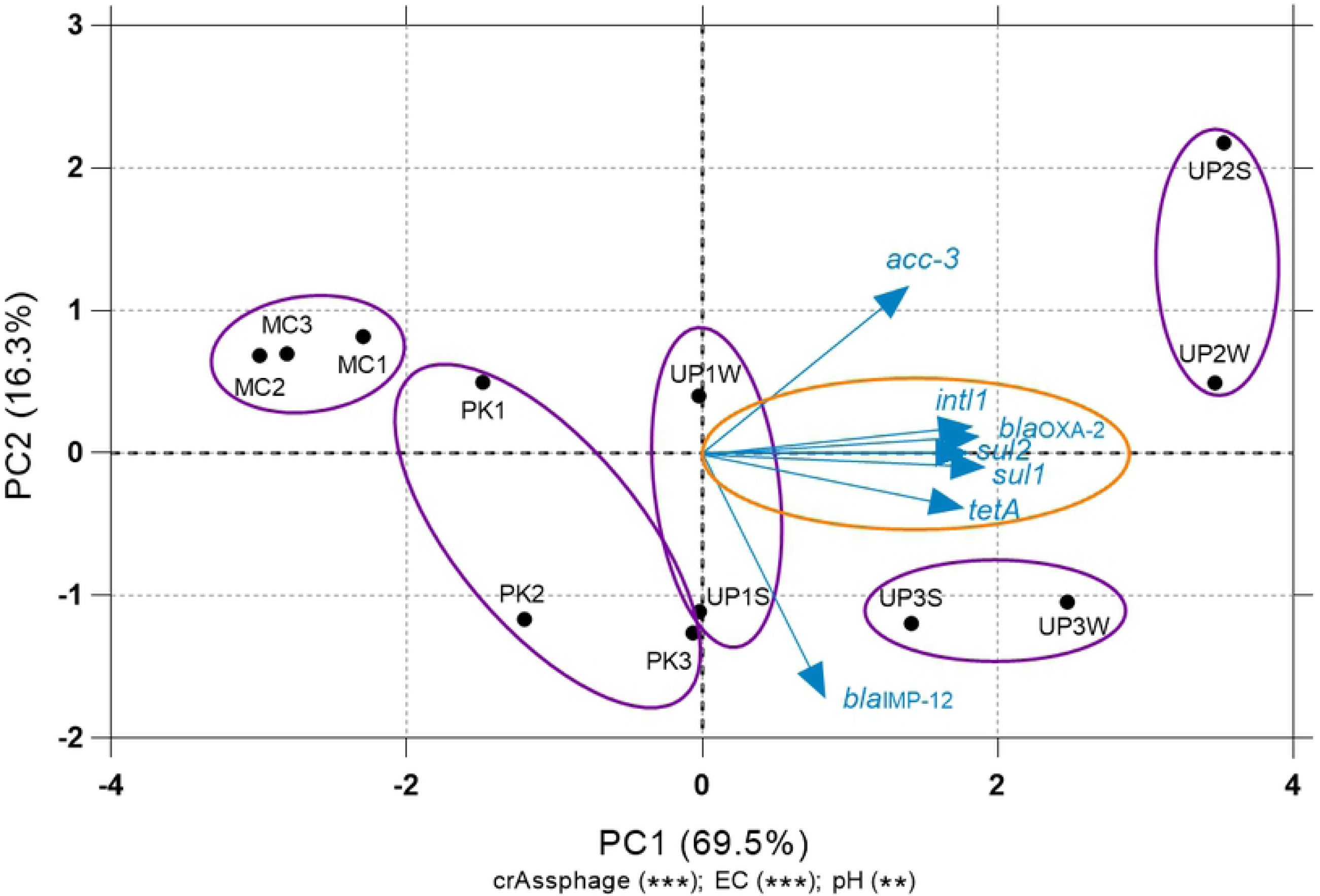
Principal Component Analysis (PCA) and Multiple Regression. The PCA was performed with the data of normalized gene abundance per site of ARGs and *intl1*. The first two PCA axis were recovered and used to perform a multiple linear regression (Lm) of the PCs with respect of the levels of metals, pH, EC, and the normalized abundance of the fecal pollution marker crAssphage. The parameters for the models were automatically selected in function of its R^2^ adjusted value and its variance inflation factor (VIF) to avoid multicollinearity among environmental variables. The number in parenthesis represent the percentage of variation explained by the axis, and the parameters that are significantly related with the axis are represented with: * p<0.05, ** p<0.001 and ***p<0.0001. Related sampling points are indicated within purple circles. The abundance of *intl1* and the ARGs are represented as blue arrows, the direction of the arrow indicates increasing abundance of the genes. The angle of the arrows with respect to the axis represent the linear relation of the abundance with the PC, the orange circle is showing the most important variables (*intl1, sul1, sul2, bla*_OXA-2_, *tetA*) which show the lowest angles of their corresponding arrows with respect to PC1. Along the PC1 the abundance of *intl1* and ARGs increase toward the right. S: sediments; W: water.

## Discussion

To our knowledge, the present study is the first to explore the relationship between ARGs, metal pollution, and wastewater discharges. In order to establish these relationships, we analyzed the abundance of different ARGs and the class 1 integron (*intl1*) in three water bodies. PK site, is a glacier lake which could be considered an ecological intact environment with very few anthropogenic activities around. Like PK, MC, is also a glacier lake but heavily impacted by mining discharges. UP, the third site, is a peri-urban lake with a long history of receiving both mining and wastewater discharges. Heavy metal levels were measured in sediment samples and we also quantified the human fecal pollution marker crAssphage along with the abundance of different ARGs. Our results suggest that fecal contribution was the major driver of increased abundance of *intl1* and ARGs included in this study. This conclusion offers a different scenario from what has been suggested in other studies that favor metals as selective agents of microbial antibiotic resistance (12-15), through co-selection process in the environment (11). However, it is important to note that fecal pollution was not considered in any of these studies. On the other hand, our results are in good agreement with a recent study that concluded ARGs abundance, in almost all cases, can be explained by the human fecal contributions in human impacted environments (18). Further analysis, especially metagenomic approaches, are needed to clarify whether the abundance of ARGs, including those not detected in our analysis, is related to fecal pollution or metal contamination in water bodies impacted by both, mining activities and wastewater.

The absence of crAssphage suggests that PK is a pristine lake, while MC could be considered an extreme environment. MC samples presented very low pH, high EC and elevated metal levels (Fig 2) characteristics associated with AMD impacted sites (22). These features were expected as mining activities at large scale were performed in the Milluni area until 1990; since then, only small cooperatives operate in the valley (38). Our results support previous studies that indicate AMD discharges on surface water acidified and enriched metal (As, Fe, Pb, Cd, Zn, Cu, Sn) dissolution and mobility (26). Therefore, MC represents a very harsh environment for microbial life. Remarkably we were able to detect and quantify 4 out of 14 ARGs on this site. Whether these genes are functional and which species contain them is a topic that must be investigated in future studies. The Pata Khota lake, on the other hand, presented low metal levels, almost neutral pH and low EC, as expected for an intact ecological environment. The chemical composition of PK is characteristic of natural lakes at high altitudes in the mountains (27). The mineralogical composition explains the levels of metals found in this pristine site (26). The Milluni valley has little anthropogenic impact other than mining and camelid cattle raising for subsistence. Although, it has been shown that the crAssphage gene that we used can be found in water with animal fecal content (20), crAssphage could not be detected in Pata Khota and Milluni Chico (Fig 4). The latter could be explained because of the extreme conditions that may influence the survival of crAssphage host in water. In the case of Pata Khota, the absence of this fecal pollution marker could reflect either, the absence of its host or the pristine character of this site.

In agreement with previous studies (28), Uru Uru lake was characterized by high EC and alkaline pH. UP3 presented the highest levels of metals among all points, followed by MC sampling points. UP1, a point that receives water from an old open pit gold mine transformed into an artificial meromictic lake presented lower values of metals compared with MC points. This observation could also explain why UP1 EC values are more similar to PK, the site considered pristine. Tagarete channel (UP2) metal levels were very low and similar to those of PK lake. UP2 receives wastewater discharges from Oruro city and disembogues in the Uru Uru lake (UP3). Consistent with this fecal pollution input, all Uru Uru samples (except UP1 sediments) were positive for the crAssphage marker. Therefore, Uru Uru sampling sites are simultaneously impacted by both fecal pollution and mining discharges.

We analyzed the abundance of fourteen ARGs, and the mobile genetic element *intl1.* Overall seven ARGs were detected in our samples, and *Intl1* was detected in all of them. The most abundant genes were *intl1, bla*_OXA-2_, *sul1* and *sul2*. All these genes presented positive correlations among them (Fig 5). Commonly *intl1* can be found in the environment positively correlated with several ARGs (6) and its abundance is strongly correlated with the abundance of multi-drug resistant bacteria (39). *intl1* and *sul1* are located together on MGEs and hence linked (40, 41). Previously, another study reported the presence of *sul1* and *sul2* in the Milluni valley, specifically in Pata Khota. When they analyzed the levels of sulfamethoxazole in water, the antibiotic was not detected (32) suggesting that the presence of these genes can occur naturally in bacteria residing in these aquatic environments, as has been found in other pristine sites (42-44). Taking into consideration that PK and MC have little antropogenic activity around, and that antibiotic levels are reported undetectable in one of our study sites (32) we assume antibiotics will not play a major rol in our analysis. Although we did not measured antibiotic levels inour sampled sites, erythromycin levels were reported under the limit of detection or in the order of ng/g of sediment for UP1 and UP2 nearby sites, respectively (Guzman-Otazo *et al*. In preparation). Even if other antibiotics could be present, their levels would be expected to be residual, given the dilution effect that they encounter in these large water bodies. Furthermore, recent evidence suggest that antibiotic residual concentrations do not play an important role in ARGs abundance (18).

Some studies showed that metals such as Cu, Zn, Cd, and Ni can exert stronger selection pressure over environmental microbial communities favoring the selection of resistant bacteria, even more than antibiotics themselves (13, 45). Even though metal levels in MC were higher than in PK, we found similar ARGs abundance in both lakes. In fact, they clustered together according to its ARGs abundance (Fig 4) and were very close to each other in the PCA analysis (Fig 7). Samples collected in UP2, that drains sewage from Oruro city to UP3 presented the highest abundance of ARGs and *intl1*. Remarkably, there are significantly higher levels of all the measured metals in UP3. However, the ARGs abundance in both sites grouped together in the hierarchical clustering analysis (Fig 4). These results suggest that other parameters different from metal levels are explaining the variation in ARGs abundance.

Previous studies reported positive correlations of ARGs and crAssphage (18, 46, 47). The hierarquical clustering analysis (Fig 4) revealed that higher levels of crAssphage grouped most of the UP2 and UP3 samples, which at the same time presented the highest levels of six out of the seven quantified genes. Also, we showed that samples with and without fecal pollution presented statistically significant differences in the abundance of *intl1, sul1, sul2, bla*_OXA-2_ and *tetA* (Fig 6). All these genes presented a positive strong correlation among them, but no individual correlation between the abundance of crAssphage and the other genes were found (Fig 5). These findings could be explained by the low number of crAssphage positive samples (only five). Moreover, the Tagarete channel strongly impacted by fecal discharges was the only point in which *acc-3* was detected, suggesting that the accumulation of bacteria from feces might be the main source of ARGs. Other studies used crAssphage as a molecular marker able to track human fecal pollution in aquatic environments (20, 48). Our results support the use of crAssphage as a marker for human fecal pollution and as a proxy to predict the presence of ARGs in wastewater impacted aquatic environments (46, 47).

Our PCA results on *intl1* and ARGs abundance showed that PC1 (69.5% of the variation) had a strong linear relationship with *Intl1, sul1, sul2, bla*_OXA-2_ and *tetA*. A multiparametric linear model revealed that the most important factors explaining this variation in this axis were EC, neutral pH, and crAssphage abundance. EC is an indicator of anthropogenic impact as values increase with mining and sewage discharges (22, 49). It is well known that pH is considered the most important factor influencing the microbial community composition in soils (50, 51), and microbial community composition is the most important factor determining the resistome in soils at continental levels (52). Therefore, it is possible that both pH and EC are indirectly conditioning the resistome in mining and wastewater impacted environments. It is important to note that pH values were only measured in water of each sampling site. Although overlying waters do not necessarily correlate with the pH of sediments in acidified environments (53), previous studies on the Milluni basin (26) reported similar sediment pH values.

Taken together our results suggest that likely fecal pollution but not metal contamination better explains the abundance of ARGs associated with *intl1* in aquatic environments impacted by both, mining and wastewater discharges. Other important factors explaining the abundance of ARGs are the physicochemical conditions (pH and EC), which can determine the composition of microbial communities and, thus, the resistome in these environments.

## Acknowledgements

The authors thank Dario Acha for help with the metal analysis and Nataniel Mamani for his technical assistance. This study was supported by grants from the Swedish Research Council to ÅS, EJ, SG-C and CC (dnr 2017-05423 and 2019-04202), and a bolivian grant from Impuesto Directo a los Hidrocarburos to SG-C (IDH 2013-2014).

## References

1. Llor C, Bjerrum L. Antimicrobial resistance: risk associated with antibiotic overuse and initiatives to reduce the problem. Therapeutic Advances in Drug Safety. 2014;5(6):229–41.

2. Berendonk TU, Manaia CM, Merlin C, Fatta-Kassinos D, Cytryn E, Walsh F, et al. Tackling antibiotic resistance: the environmental framework. Nature reviews Microbiology. 2015;13(5):310–7.

3. Gillings MR, Westoby M, Ghaly TM. Pollutants That Replicate: Xenogenetic DNAs. Trends Microbiol. 2018.

4. Marti E, Variatza E, Balcazar JL. The role of aquatic ecosystems as reservoirs of antibiotic resistance. Trends Microbiol. 2014;22(1):36–41.

5. Chu BTT, Petrovich ML, Chaudhary A, Wright D, Murphy B, Wells G, et al. Metagenomics Reveals the Impact of Wastewater Treatment Plants on the Dispersal of Microorganisms and Genes in Aquatic Sediments. Applied and Environmental Microbiology. 2018;84(5).

6. Zhu Y-G, Zhao Y, Li B, Huang C-L, Zhang S-Y, Yu S, et al. Continental-scale pollution of estuaries with antibiotic resistance genes. Nature Microbiology. 2017;2:16270.

7. Jutkina J, Marathe NP, Flach CF, Larsson DGJ. Antibiotics and common antibacterial biocides stimulate horizontal transfer of resistance at low concentrations. Science of The Total Environment. 2018;616-617:172–8.

8. Nesme J, Cecillon S, Delmont TO, Monier JM, Vogel TM, Simonet P. Large-scale metagenomic-based study of antibiotic resistance in the environment. Curr Biol. 2014;24(10):1096–100.

9. Navarro MC, Pérez-Sirvent C, Martínez-Sánchez MJ, Vidal J, Tovar PJ, Bech J. Abandoned mine sites as a source of contamination by heavy metals: A case study in a semi-arid zone. Journal of Geochemical Exploration. 2008;96(2):183–93.

10. Liu J, Yin M, Luo X, Xiao T, Wu Z, Li N, et al. The mobility of thallium in sediments and source apportionment by lead isotopes. Chemosphere. 2019;219:864–74.

11. Baker-Austin C, Wright MS, Stepanauskas R, McArthur JV. Co-selection of antibiotic and metal resistance. Trends in Microbiology. 2006;14(4):176–82.

12. Zhao Y, Cocerva T, Cox S, Tardif S, Su J-Q, Zhu Y-G, et al. Evidence for co-selection of antibiotic resistance genes and mobile genetic elements in metal polluted urban soils. Science of The Total Environment. 2019;656:512–20.

13. Song J, Rensing C, Holm PE, Virta M, Brandt KK. Comparison of Metals and Tetracycline as Selective Agents for Development of Tetracycline Resistant Bacterial Communities in Agricultural Soil. Environ Sci Technol. 2017;51(5):3040–7.

14. Zhao Z, Wang J, Han Y, Chen J, Liu G, Lu H, et al. Nutrients, heavy metals and microbial communities co-driven distribution of antibiotic resistance genes in adjacent environment of mariculture. Environmental Pollution. 2016.

15. Gao P, He S, Huang S, Li K, Liu Z, Xue G, et al. Impacts of coexisting antibiotics, antibacterial residues, and heavy metals on the occurrence of erythromycin resistance genes in urban wastewater. Appl Microbiol Biotechnol. 2015;99(9):3971–80.

16. Taylor NG, Verner-Jeffreys DW, Baker-Austin C. Aquatic systems: maintaining, mixing and mobilising antimicrobial resistance? Trends Ecol Evol. 2011;26(6):278–84.

17. Baquero F, Martínez J-L, Cantón R. Antibiotics and antibiotic resistance in water environments. Current Opinion in Biotechnology. 2008;19(3):260–5.

18. Karkman A, Pärnänen K, Larsson DGJ. Fecal pollution can explain antibiotic resistance gene abundances in anthropogenically impacted environments. Nature communications. 2019;10(1):80.

19. Dutilh BE, Cassman N, McNair K, Sanchez SE, Silva GG, Boling L, et al. A highly abundant bacteriophage discovered in the unknown sequences of human faecal metagenomes. Nature communications. 2014;5:4498.

20. García-Aljaro C, Ballesté E, Muniesa M, Jofre J. Determination of crAssphage in water samples and applicability for tracking human faecal pollution. Microbial biotechnology. 2017;10(6):1775–80.

21. Huang X, Sillanpää M, Gjessing ET, Peräniemi S, Vogt RD. Environmental impact of mining activities on the surface water quality in Tibet: Gyama valley. Science of The Total Environment. 2010;408(19):4177–84.

22. Gray NF. Field assessment of acid mine drainage contamination in surface and ground water. Environmental Geology. 1996;27(4):358–61.

23. Satgé F, Hussain Y, Xavier A, Zolá RP, Salles L, Timouk F, et al. Unraveling the impacts of droughts and agricultural intensification on the Altiplano water resources. Agricultural and Forest Meteorology. 2019;279:107710.

24. Poma V, Mamani N, Iñiguez V. Impact of urban contamination of the La Paz River basin on thermotolerant coliform density and occurrence of multiple antibiotic resistant enteric pathogens in river water, irrigated soil and fresh vegetables. SpringerPlus. 2016;5(1).

25. Guzman-Otazo J, Gonzales-Siles L, Poma V, Bengtsson-Palme J, Thorell K, Flach CF, et al. Diarrheal bacterial pathogens and multi-resistant enterobacteria in the Choqueyapu River in La Paz, Bolivia. PLoS One. 2019;14(1):e0210735.

26. Salvarredy-Aranguren MM, Probst A, Roulet M, Isaure M-P. Contamination of surface waters by mining wastes in the Milluni Valley (Cordillera Real, Bolivia): Mineralogical and hydrological influences. Applied Geochemistry. 2008;23(5):1299–324.

27. Driesen K. Contamination of surface waters by the former mining industry in the Milluni Valley (Cordillera Real, Bolivia) and the application of the water planning model WEAP. Bélgica: University College Ghent; 2012.

28. Alanoca L, Guedron S, Amouroux D, Audry S, Monperrus M, Tessier E, et al. Synergistic effects of mining and urban effluents on the level and distribution of methylmercury in a shallow aquatic ecosystem of the Bolivian Altiplano. Environ Sci Process Impacts. 2016;18(12):1550–60.

29. Zamora Echenique G, Mata Parello J, Quezada H R. Propuesta de “Geoparque Oruro – Bolivia” Alternativa para la preservación del patrimonio geológico, minero y sociocultural. REV MAMYM. 2018(5):19 – 33.

30. Guedron S, Point D, Acha D, Bouchet S, Baya PA, Tessier E, et al. Mercury contamination level and speciation inventory in Lakes Titicaca & Uru-Uru (Bolivia): Current status and future trends. Environmental pollution (Barking, Essex : 1987). 2017;231(Pt 1):262–70.

31. Fang Y, Xu M, Chen X, Sun G, Guo J, Wu W, et al. Modified pretreatment method for total microbial DNA extraction from contaminated river sediment. Frontiers of Environmental Science & Engineering. 2015;9(3):444–52.

32. Archundia D, Duwig C, Lehembre F, Chiron S, Morel MC, Prado B, et al. Antibiotic pollution in the Katari subcatchment of the Titicaca Lake: Major transformation products and occurrence of resistance genes. Sci Total Environ. 2017;576:671–82.

33. Alcock BP, Raphenya AR, Lau TTY, Tsang KK, Bouchard M, Edalatmand A, et al. CARD 2020: antibiotic resistome surveillance with the comprehensive antibiotic resistance database. Nucleic Acids Res. 2020;48(D1):D517–d25.

34. Ye J, Coulouris G, Zaretskaya I, Cutcutache I, Rozen S, Madden TL. Primer-BLAST: a tool to design target-specific primers for polymerase chain reaction. BMC Bioinformatics. 2012;13:134.

35. Nolvak H, Truu M, Truu J. Evaluation of quantitative real-time PCR workflow modifications on 16S rRNA and tetA gene quantification in environmental samples. Sci Total Environ. 2012;426:351–8.

36. Zhu Y-G, Johnson TA, Su J-Q, Qiao M, Guo G-X, Stedtfeld RD, et al. Diverse and abundant antibiotic resistance genes in Chinese swine farms. Proceedings of the National Academy of Sciences. 2013;110(9):3435–40.

37. Zhang Z, Qu Y, Li S, Feng K, Wang S, Cai W, et al. Soil bacterial quantification approaches coupling with relative abundances reflecting the changes of taxa. Scientific reports. 2017;7(1):4837.

38. Rios C. Estudio de la Contaminación Ambiental por las Descargas Mineras de Comsur en la Represa Milluni. La Paz, Bolivia.: Universidad Mayor de San Andres; 1985.

39. Lucassen R, Rehberg L, Heyden M, Bockmühl D. Strong correlation of total phenotypic resistance of samples from household environments and the prevalence of class 1 integrons suggests for the use of the relative prevalence of intI1 as a screening tool for multi-resistance. PLOS ONE. 2019;14(6):e0218277.

40. Uyaguari-Díaz MI, Croxen MA, Luo Z, Cronin KI, Chan M, Baticados WN, et al. Human Activity Determines the Presence of Integron-Associated and Antibiotic Resistance Genes in Southwestern British Columbia. Frontiers in microbiology. 2018;9:852-.

41. Gillings MR, Gaze WH, Pruden A, Smalla K, Tiedje JM, Zhu Y-G. Using the class 1 integron-integrase gene as a proxy for anthropogenic pollution. The ISME journal. 2015;9(6):1269–79.

42. Storteboom H, Arabi M, Davis JG, Crimi B, Pruden A. Tracking Antibiotic Resistance Genes in the South Platte River Basin Using Molecular Signatures of Urban, Agricultural, And Pristine Sources. Environmental Science & Technology. 2010;44(19):7397–404.

43. Ouyang W-Y, Huang F-Y, Zhao Y, Li H, Su J-Q. Increased levels of antibiotic resistance in urban stream of Jiulongjiang River, China. Applied Microbiology and Biotechnology. 2015;99(13):5697–707.

44. Van Goethem MW, Pierneef R, Bezuidt OKI, Van De Peer Y, Cowan DA, Makhalanyane TP. A reservoir of ‘historical’ antibiotic resistance genes in remote pristine Antarctic soils. Microbiome. 2018;6(1):40.

45. Stepanauskas R, Glenn TC, Jagoe CH, Tuckfield RC, Lindell AH, King CJ, et al. Coselection for microbial resistance to metals and antibiotics in freshwater microcosms. Environmental Microbiology. 2006;8(9):1510–4.

46. Stachler E, Crank K, Bibby K. Co-Occurrence of crAssphage with Antibiotic Resistance Genes in an Impacted Urban Watershed. Environmental Science & Technology Letters. 2019;6(4):216–21.

47. Chen H, Bai X, Li Y, Jing L, Chen R, Teng Y. Source identification of antibiotic resistance genes in a peri-urban river using novel crAssphage marker genes and metagenomic signatures. Water research. 2019;167:115098.

48. Stachler E, Akyon B, de Carvalho NA, Ference C, Bibby K. Correlation of crAssphage qPCR Markers with Culturable and Molecular Indicators of Human Fecal Pollution in an Impacted Urban Watershed. Environmental Science & Technology. 2018;52(13):7505–12.

49. Pal M, Samal N, Roy P, Biswas Roy M. Electrical Conductivity of Lake Water as Environmental Monitoring –A Case study of Rudra sagar Lake. Journal of Environmental Science, Toxicology and Food Technology. 2015.

50. Lauber CL, Hamady M, Knight R, Fierer N. Pyrosequencing-Based Assessment of Soil pH as a Predictor of Soil Bacterial Community Structure at the Continental Scale. Applied and Environmental Microbiology. 2009;75(15):5111–20.

51. Bahram M, Hildebrand F, Forslund SK, Anderson JL, Soudzilovskaia NA, Bodegom PM, et al. Structure and function of the global topsoil microbiome. Nature. 2018;560(7717):233–7.

52. Forsberg KJ, Patel S, Gibson MK, Lauber CL, Knight R, Fierer N, et al. Bacterial phylogeny structures soil resistomes across habitats. Nature. 2014;509(7502):612–6.

53. Herlihy AT, Mills AL. The pH regime of sediments underlying acidified waters. Biogeochemistry. 1986;2(1):95–9.

